# Histone H3 T11 phosphorylation by Sch9 and CK2 regulates lifespan by controlling the nutritional stress response

**DOI:** 10.1101/282384

**Authors:** Seunghee Oh, Tamaki Suganuma, Madelaine M. Gogol, Jerry L. Workman

## Abstract

Upon nutritional stress, the metabolic status of cells is changed by nutrient signaling pathways to ensure survival. Altered metabolism by nutrient signaling pathways has been suggested to influence cellular lifespan. However, it remains unclear how chromatin regulation is involved in this process. Here, we found that histone H3 threonine 11 phosphorylation (H3pT11) functions as a marker for nutritional stress and aging. Sch9 and CK2 kinases cooperatively regulate H3pT11 under stress conditions. Importantly, H3pT11 defective mutants prolonged chronological lifespan by altering nutritional stress responses. Thus, the phosphorylation of H3T11 by Sch9 and CK2 engages a nutritional stress response to chromatin in the regulation of lifespan.

## Introduction

Nutritional stress is an unavoidable event for most of organisms, and appropriate metabolic adaptation to nutritional deficiency is essential to ensure the survival of cells and organisms. Given the fact that calorie restriction relates to the regulation of lifespan from yeast to mammals (Guarente, 2006), proper adaptations of metabolism due to nutritional changes may be critical processes in the regulation of lifespan. *Saccharomyces cerevisiae* utilizes different carbon sources and adapts to various nutritional environments by changing its metabolism (Broach, 2012). In yeast, glucose is the most preferred carbon source for their growth. When external glucose levels are sufficient, they utilize fermentation for energy production even if the oxygen concentration is high, which resembles the Warburg effect seen in stem cells and cancer cells (Vander Heiden, Cantley, & Thompson, 2009). When the levels of glucose and other fermentable carbon source run low, they shift energy metabolism from fermentation to the mitochondrial respiration pathway. Multiple signaling pathways including PKA/Ras, TOR, Sch9 cooperate to regulate the metabolic transition (Galdieri, Mehrotra, Yu, & Vancura, 2010; Wilson & Roach, 2002), which is accompanied by global changes in gene expression (DeRisi, Iyer, & Brown, 1997). Many factors important for regulation of the metabolic transition are also involved in the process of cellular aging (Cheng et al., 2007). Downregulation of the TOR, Sch9, and PKA/Ras2 pathways leads extension of lifespan (Fabrizio, Pozza, Pletcher, Gendron, & Longo, 2001; Longo, 1999; Powers, Kaeberlein, Caldwell, Kennedy, & Fields, 2006; Wei et al., 2008).

Chromatin modifying enzymes also play roles in aging (Benayoun, Pollina, & Brunet, 2015; Sen, Shah, Nativio, & Berger, 2016). The sirtuin deacetylase Sir2 regulates lifespan by reducing histone H4 lysine 16 acetylation (H4K16ac) levels at telomere (W. Dang et al., 2009). Inactivation of a chromatin remodeling protein, Isw1, extends lifespan by induction of genotoxic stress response genes (Weiwei Dang et al., 2014). However, direct connections between nutrition sensing pathways and chromatin regulation in the aging process are still unknown. Interestingly, pyruvate kinases in yeast and humans have been shown to phosphorylate H3 at T11 (Li et al., 2015; Yang et al., 2012), suggesting that H3pT11 mediates a connection between metabolism and chromatin. Several different kinases are responsible for H3T11 phosphorylation. In yeast, Mek1 directly regulate H3T11 phosphorylation during meiosis (Govin et al., 2010; Kniewel et al., 2017). In human, protein kinase N1, PKN1, phosphorylates H3T11 at promoters of androgen receptor dependent genes (Metzger et al., 2008), and checkpoint kinase 1, Chk1, phosphorylates H3T11 in mouse embryonic fibroblast cells (Shimada et al., 2008).

Casein kinase 2 complex, CK2, is a ubiquitous serine/threonine kinase complex and plays roles in cell growth and proliferation. CK2 is a conserved protein complex from yeast to human. Yeast CK2 consists of two catalytic subunits (a1 and a2) and two regulatory subunits (b1 and b2) (Ahmed, Gerber, & Cochet, 2002; Litchfield, 2003). CK2 phosphorylates many kinds of substrates including histones (Basnet et al., 2014; Cheung et al., 2005; Franchin et al., 2017), and this pleiotropy implies a broad function of CK2 in various biological pathways including glucose metabolism (Borgo et al., 2017).

Interestingly, deletion of a CK2 catalytic subunit, Cka2, extends lifespan in yeast; however, the mechanism of how CK2 regulates lifespan is unknown (Fabrizio et al., 2010).

Here, we found that upon nutritional stress in yeast, the level of H3T11 phosphorylation specifically increased at stress responsive genes and regulates transcription of genes involved the metabolic transition. We also found that Sch9 and Cka1, a catalytic subunit of CK2, are required for the phosphorylation of H3pT11 under the stress. Importantly, loss of H3T11 phosphorylation prolongs lifespan by altering the stress response at an early stage of chronological lifespan (CLS), suggesting that H3T11 phosphorylation by CK2 and Sch9 links nutritional stress to chromatin during the process of aging.

## Results

### H3T11 phosphorylation is increased upon the nutritional stress

Our previous study implied a connection between H3pT11 and glucose metabolism (Li et al., 2015), we therefore examined the relationship between H3pT11 and external glucose levels using an antibody specific to H3pT11 (validated in Figure 1-figure supplement 1). Culture media for wild type (WT) cells was changed from glucose rich (2%) YPD to YP with different concentrations of glucose (0.02%, 0.2%, or 2%). The global levels of H3pT11 showed a clear negative correlation with media glucose levels (Figure 1A). When the culture media was shifted from YPD to YP with 3% glycerol (YPglycerol), which is non-fermentable and is nutritionally unfavorable carbon source for yeast, we also observed robust increases in the levels of H3pT11 (Figure 1B). H3pT11 levels were not changed in media containing both glucose and glycerol compared to the 2% glucose condition, and direct addition of glucose was sufficient to suppress the increase of H3pT11 levels found in YPglycerol (Figure 1-figure supplement 2A). These data demonstrate that H3pT11 levels are specifically increased in low glucose conditions.

**Figure 1.**
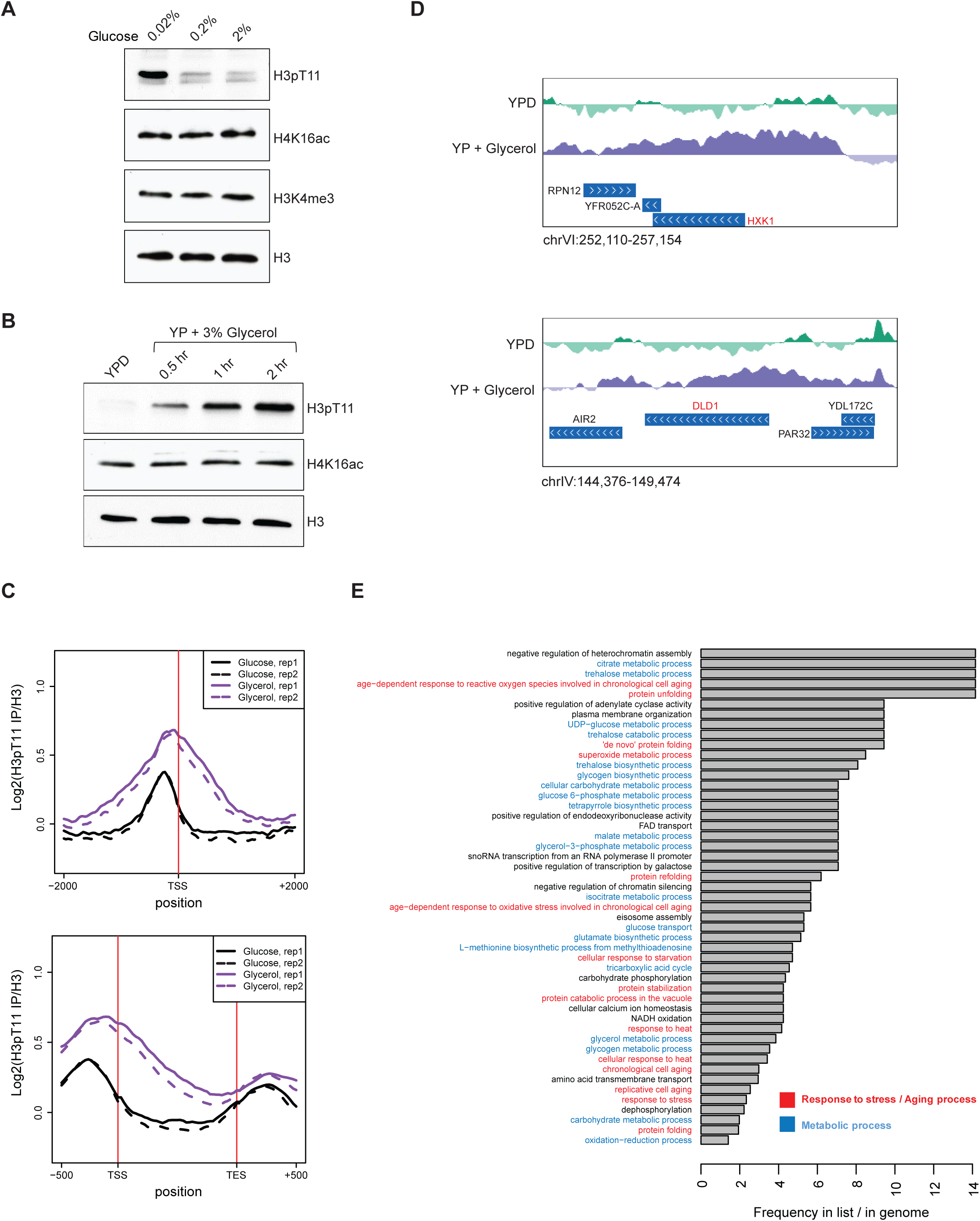
H3pT11 responds to nutritional stress. (A and B) H3pT11 levels in different concentration of glucose (A) or non-fermentable glycerol (B) containing media conditions measured by Western blots. (C) The average profiles of H3pT11 at 366 genes, whose H3pT11 levels are increased in YPglycerol (Glycerol) compared to YPD (Glucose) condition. TSS, transcription start site; TES, transcription end site. (D) Normalized H3pT11 levels to H3 at *HXK1* and *DLD1* gene loci in YPD and YPglycerol conditions. (E) GO term analysis of the 366 genes shown in (C).

To ask how the genomic distribution of H3pT11 changes upon nutritional stress, we performed ChIP-sequencing (ChIP-seq) of H3pT11 in cells cultured in YPD or YPglycerol conditions. In agreement with the Western blots (Figure 1B), the total number of H3pT11 peaks increased in YPglycerol conditions compared to YPD (Figure 1-figure supplement 2B). We identified 366 genes, whose H3pT11 levels were increased upon this nutritional stress (Figure 1C). These genes included hexokinase, *HXK1*, and mitochondrial lactate dehydrogenase, *DLD1*, whose expression are known to increase in low glucose conditions (Lodi, Alberti, Guiard, & Ferrero, 1999; Rodriguez, De La Cera, Herrero, & Moreno, 2001) (Figure 1D). Through GO term analysis of the genes, where H3pT11 levels were changed upon the stress (YPglycerol), we found that the genes with increased H3pT11 levels were highly enriched in aging-related processes, stress responses, and metabolic pathways (Figure 1E). H3pT11 levels were decreased at 139 genes (Figure 1-figure supplement 2C) in YPglycerol compared to YPD. The genes with reduced H3pT11 levels were involved in fermentation and translation, which are generally repressed in nutritional stress conditions (Figure 1-figure supplement 2D). These results show that H3pT11 levels are specifically changed at a group of genes involved in the nutritional stress responses.

### H3T11 phosphorylation regulates transcription upon nutritional stress

The genome-wide distribution of H3pT11 strongly suggests that H3pT11 has a role in regulation of the transcriptional response to nutritional stress. We classified RNA polymerase II (Pol II) regulated genes into 5 groups based on their mRNA expression levels of RNA-sequencing (RNA-seq) in YPglycerol condition and compared H3pT11 levels among those 5 groups. H3pT11 levels were mostly enriched in promoter regions. In these regions, the H3pT11 signals were positively correlated with mRNA expression levels in the YPglycerol condition (Figure 2A). We compared transcripts in H3T11A mutant to those in WT, cultured in YPD or YPglycerol by RNA-seq. We found a negative correlation of gene expression between YPglycerol dependence (x-axis) and H3T11A dependence (y-axis) with correlation coefficient (cor) −0.38 (Figure 2B). Thus, genes with increased expression in YPglycerol tended to be down-regulated the in H3T11A mutant, while genes with decreased expression in YPglycerol were up-regulated in H3T11A mutant.

**Figure 2.**
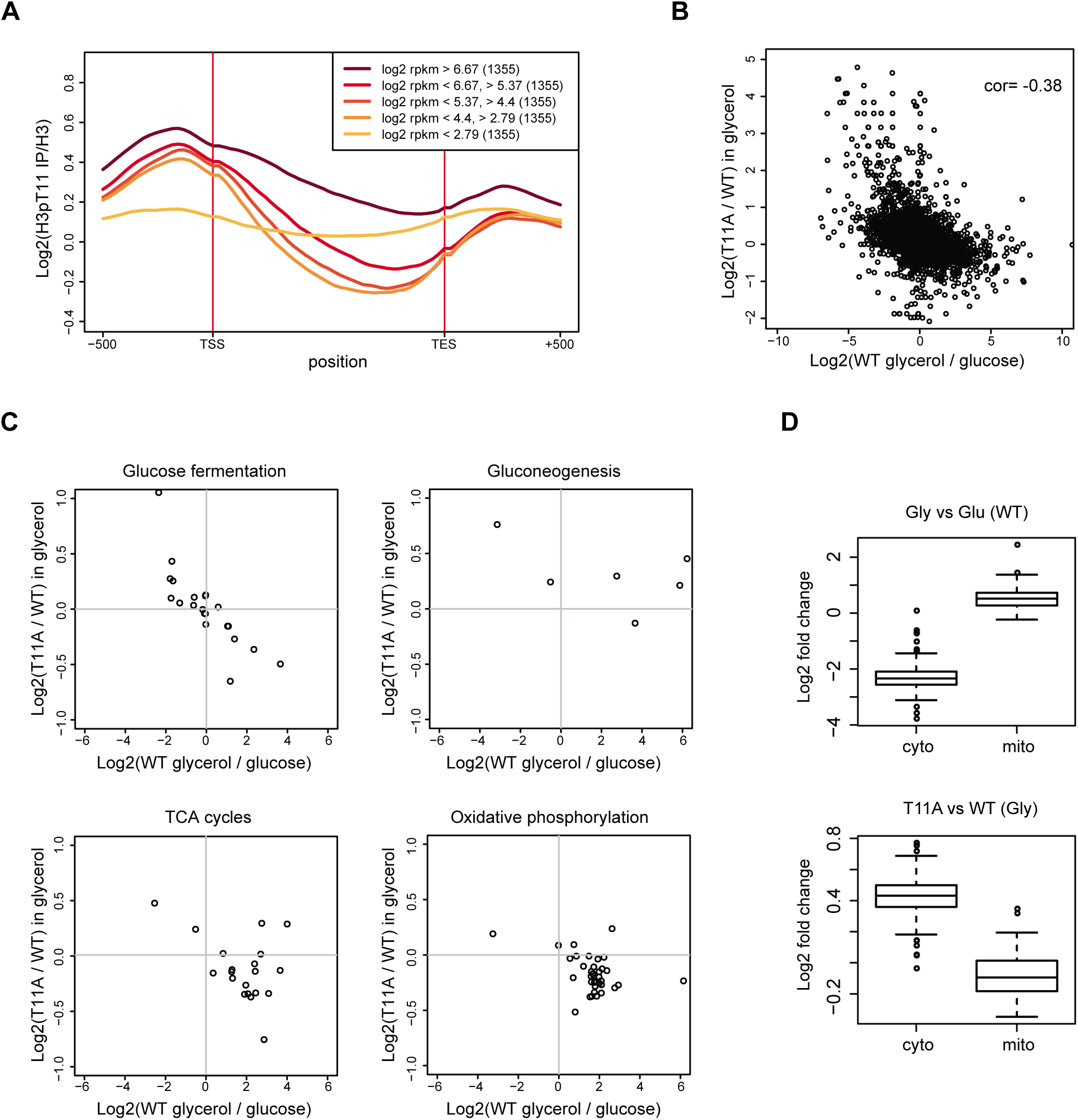
H3pT11 regulates transcription involved in metabolic transition upon nutritional stress. (A) The average H3pT11 signal of genes from different gene expression quantiles in YPglycerol. (B) A scatter plot from RNA-seq data showing a negative correlation between transcription changes upon media shift from YPD to YPglycerol (x-axis), and the changes in H3T11A mutant compared to WT in YPglycerol condition (y-axis). (C) Scatter plots from RNA-seq for transcripts of genes in indicated pathways. (D) Box-plots showing expression changes of cytoplasmic (cyto) and mitochondrial (mito) ribosomal genes upon nutritional stress condition (upper panel) and in H3T11A mutant in YPglycerol condition (lower panel). Gly, YPglycerol condition; Glu, YPD condition.

Upon nutritional stress, glucose fermentation pathway genes revealed a stronger negative correlation (cor = −0.82) between H3pT11 dependence (y-axis) and YPglycerol dependence (x-axis) compared to the correlation of all genes (cor = −0.38) (Figure 2C upper left panel). Transcripts of the genes related to the TCA cycle and the oxidative phosphorylation pathway were mostly upregulated upon the stress and relatively downregulated in the H3T11A mutant (Figure 2C lower panels). Interestingly, this trend did not match for transcripts of gluconeogenesis specific genes (Figure 2C upper right panel). The transcription of these genes was upregulated in the H3T11A mutant, regardless of their transcriptional changes in YPglycerol condition. As well as carbon source metabolism involved genes, the transcription of cellular compartment genes was also affected by H3pT11 defect. For example, there are two classes of ribosomal genes in yeast. One is cytoplasmic ribosomal genes, and the other is mitochondrial ribosomal genes. In the stress condition, the transcription of cytoplasmic ribosomal genes was downregulated, while transcription of mitochondrial ribosomal genes was upregulated (Figure 2D upper panel). Interestingly, in the H3T11A mutant, the cytoplasmic genes were upregulated, while mitochondrial genes were downregulated in YPglycerol, compared to WT (Figure 2D lower panel). Altogether, these data indicated that H3pT11 regulates the transcription of the genes involved in the metabolic transition to the mitochondrial respiratory pathways upon nutritional stress.

### Cka1 is responsible for H3T11 phosphorylation upon nutritional stress

We next asked which kinases are responsible for this modification under nutritional stress conditions. We previously showed that Pyruvate kinase 1 (Pyk1 or Cdc19) in the SESAME (<u>Se</u>rine-responsive <u>SA</u>M-containing <u>m</u>etabolic <u>e</u>nzyme) complex phosphorylates H3pT11 under nutrient rich YPD conditions (Li et al., 2015). However, Pyk1 expression is greatly reduced in low glucose conditions (Boles et al., 1997). In YPglycerol condition, we did not observe a clear difference in the global levels of H3pT11 in the SESAME subunit mutants compared to that in WT (Figure 3-figure supplement 1). This pointed to a role of different kinase(s) in H3pT11 under YPglycerol. To identify other kinase(s) responsible for H3pT11, we tested several kinases including known H3pT11 kinases in yeast and other organisms (Govin et al., 2010; Kniewel et al., 2017; Metzger et al., 2008; Shimada et al., 2008). The global levels of H3pT11 were similar among *chk1Δ*, deletion of the yeast homolog of mouse Chk1, *mek1Δ*, and WT (Figure 3-figure supplement 2A). Unexpectedly, H3pT11 was decreased in the *cka1Δ* mutant. Interestingly, H3pT11 levels were unaffected upon deletion of another catalytic subunit of CK2, *cka2Δ* mutant (Figure 3A and Figure 3-figure supplement 2A), although Cka1 and Cka2 have been thought to be functionally redundant (Chen-Wu, Padmanabha, & Glover, 1988; Padmanabha, Chen-Wu, Hanna, & Glover, 1990). Deletion of the regulatory subunits, *ckb1Δ* and *ckb2Δ*, also did not affect H3pT11 levels (Figure 3-figure supplement 2B). Thus, we examined whether CK2 complex phosphorylated H3T11 by an *in vitro* kinase assay using TAP-purified CK2 complex, recombinant H3 (rH3), and ATP. The purified CK2 complex clearly phosphorylated H3T11 (Figure 3B). We also confirmed the substrate specificity of CK2 phosphorylation at H3T11 as seen in no-signals on purified recombinant H3T11A mutant by *in vitro* kinase assay (Figure 3C). Importantly, when we measured the global levels of H3pT11 in YPglycerol condition, the H3pT11 levels were significantly reduced in *cka1Δ* mutant compared to WT or *cka2Δ* mutant (Figure 3D), indicating Cka1 is responsible for the phosphorylation of H3T11 upon nutritional stress.

**Figure 3.**
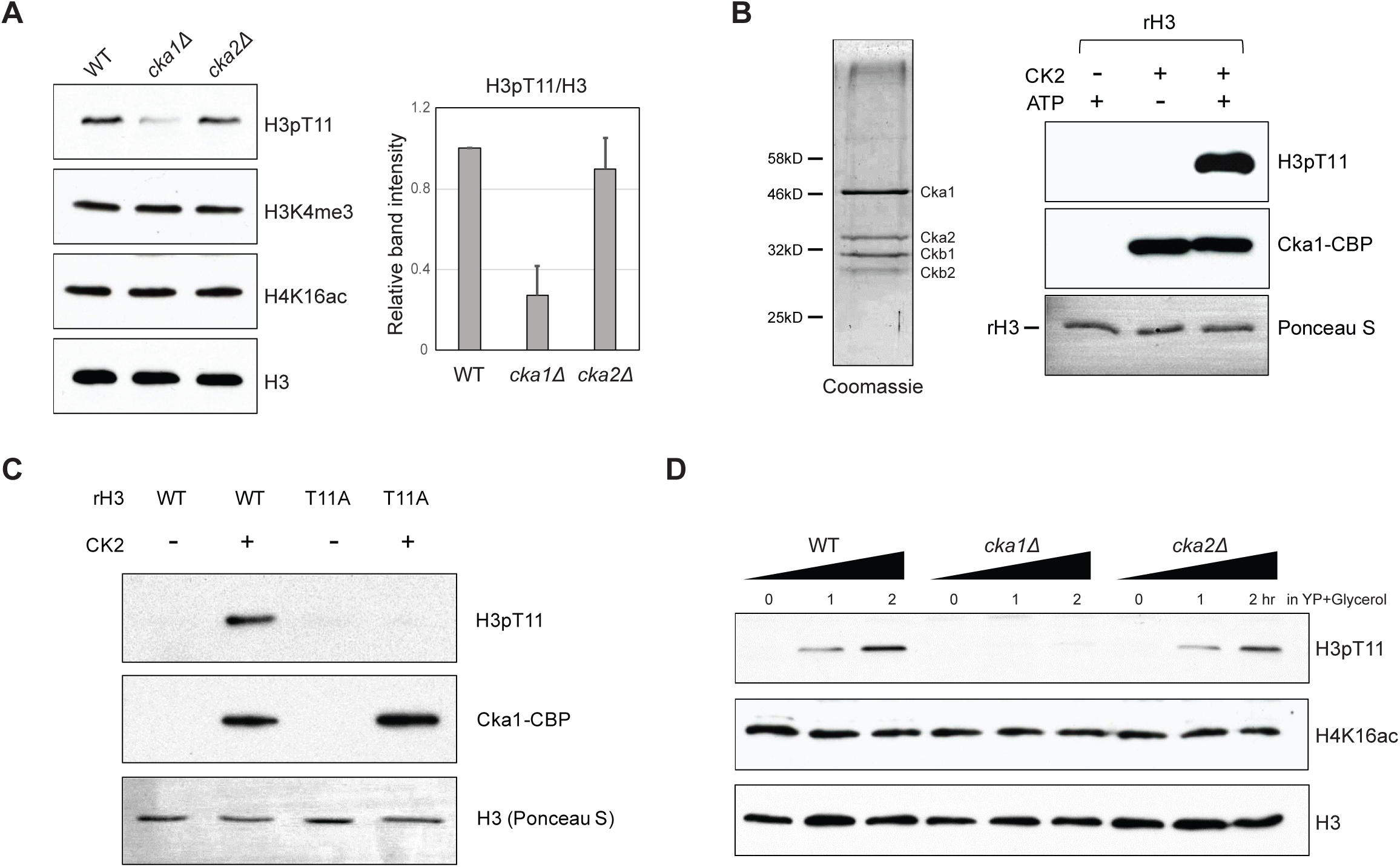
Cka1 in the CK2 complex phosphorylates H3T11. (A) (Left) H3pT11 levels of WT, *cka1Δ*, and *cka2Δ* in YPD condition analyzed by Western blots. (Right) The relative band intensities of H3pT11 to H3 signals compared to WT. Error bars represent standard deviation (STD) from three biological replicates (B) (Left) Coomassie staining of TAP purified CK2 complex using Cka1-TAP strain. (Right) *In vitro* kinase assay using TAP purified CK2 and recombinant H3 as a substrate. (C) *In vitro* kinase assay of TAP purified CK2 using recombinant H3 WT or H3T11A mutant as a substrate. (D) H3pT11 level changes in WT, *cka1Δ*, and *cka2Δ* mutants upon culture shift to YPglycerol media.

### Sch9 regulates H3T11 phosphorylation upon nutritional stress

Since H3pT11 levels respond to glucose levels in the media (Figure 1), we further examined whether H3pT11 level is related to glucose-sensing pathways. Sch9, PKA, and TOR pathways are responsible for glucose sensing in the context of calorie restriction (Powers et al., 2006). Sch9, Ras2, and Tor1 proteins are key enzymes in each pathway. To ask whether these pathways were involved in the H3pT11 regulation, we compared the H3pT11 levels among *sch9Δ, ras2Δ, tor1Δ* mutants and WT in YPglycerol condition.

Interestingly, H3pT11 increases were significantly impaired only in the *sch9Δ* mutant but not in the *ras2Δ* or *tor1Δ* mutants cultured in YPglycerol (Figure 4A), suggesting H3pT11 levels depended on Sch9 pathway. In support of this observation we found that TAP purified Sch9 protein could directly phosphorylate H3T11 *in vitro* (Figure 4B). As both CK2 and Sch9 were responsible for H3pT11 *in vivo* (Figures 3A and 4A) and were able to phosphorylate H3T11 *in vitro* (Figures 3B and 4B), we examined if CK2 and Sch9 phosphorylate H3T11 in a cooperative or independent manner *in vivo*. We measured the global levels of H3pT11 in *sch9Δcka1Δ* double mutant in YPglycerol. Interestingly, *sch9Δcka1Δ* double mutant showed similar H3pT11 level compared to *cka1Δ* alone (Figure 4C). Thus, Sch9 and CK2 are not independent of each other but have overlapping function in regulation of H3pT11 upon nutritional stress.

**Figure 4.**
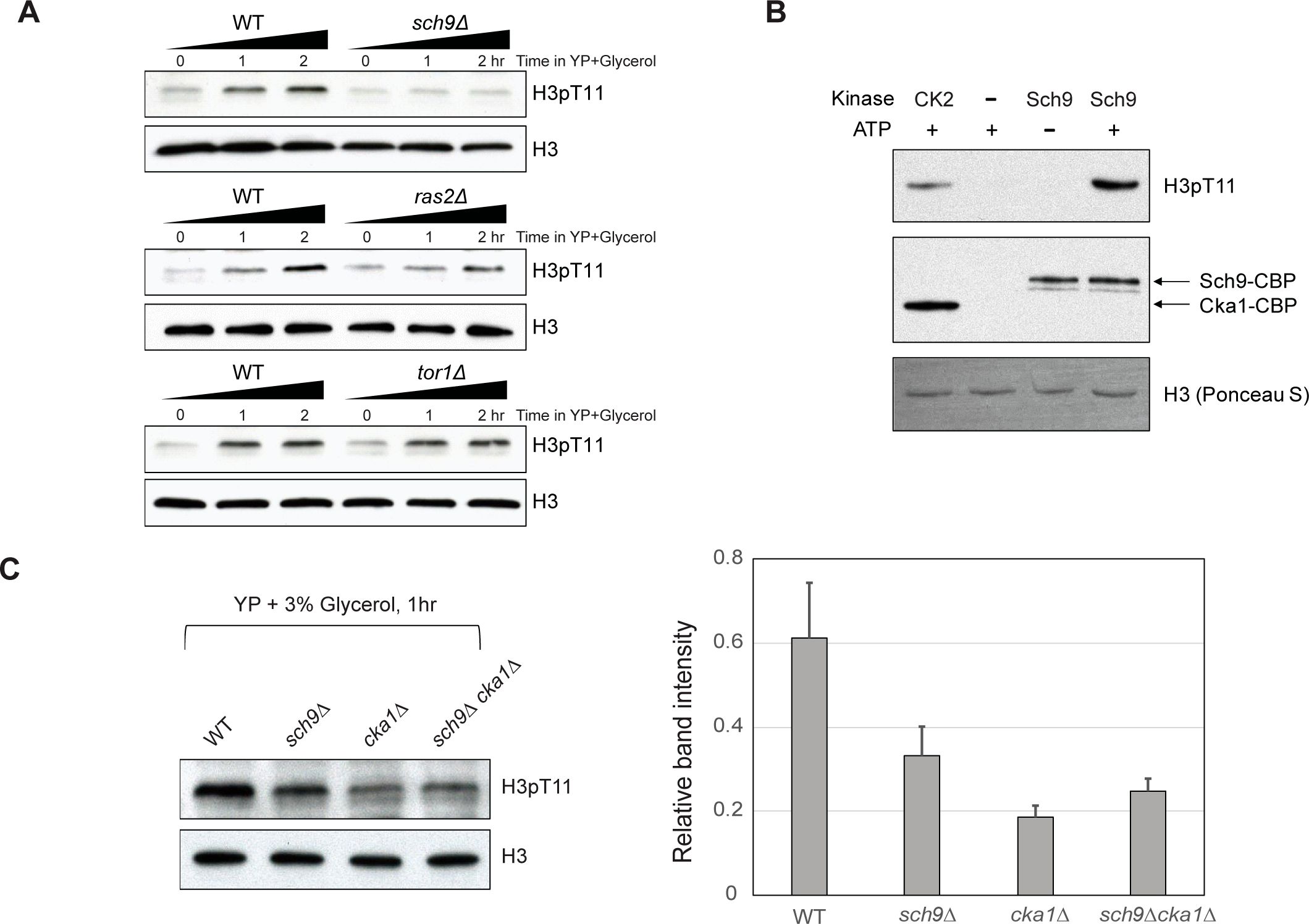
Sch9 regulates H3pT11 upon nutritional stress. (A) H3pT11 level changes of WT, *sch9Δ, ras2Δ*, and *tor1Δ* mutants upon media shift to YPglycerol measured by Western blots. (B) *In vitro* kinase assay of TAP purified (Sch9-TAP) Sch9 and CK2 using recombinant H3 as a substrate. (C) (Left) H3pT11 levels in WT, *sch9Δ, cka1Δ*, and *sch9Δcka1Δ* at 1 hour after media shift from YPD to YPglycerol analyzed by Western blots (Right) The relative ratios of H3pT11 to H3 signals are presented with error bars indicating STD from three biological replicates.

### H3T11 phosphorylation regulates lifespan

Glucose sensing pathways are closely related to lifespan control from yeast to mammals (Cheng et al., 2007), and deletion of Sch9 is a well-known long-lived mutant in yeast (Fabrizio et al., 2001). H3pT11 was tightly controlled by media glucose levels (Figure 1), and Sch9 is responsible for H3T11 phosphorylation (Figure 4). We therefore asked whether H3pT11 is involved in lifespan regulation by chronological lifespan (CLS) assays, which measures the length of time that non-dividing yeast cells survive (Longo, Shadel, Kaeberlein, & Kennedy, 2012). Strikingly, lifespan was significantly extended in the H3T11A mutant compared to the WT strain (Figure 5A). In addition, *cka1Δ* mutant also extended lifespan (Figure 5B). However, *sch9Δcka1Δ* mutant did not further extend the lifespan of *cka1Δ* or *sch9Δ* single mutant (Figure 5C). These data suggest that Sch9 and CK2 cooperatively regulate lifespan as in the case of H3pT11 regulation.

**Figure 5.**
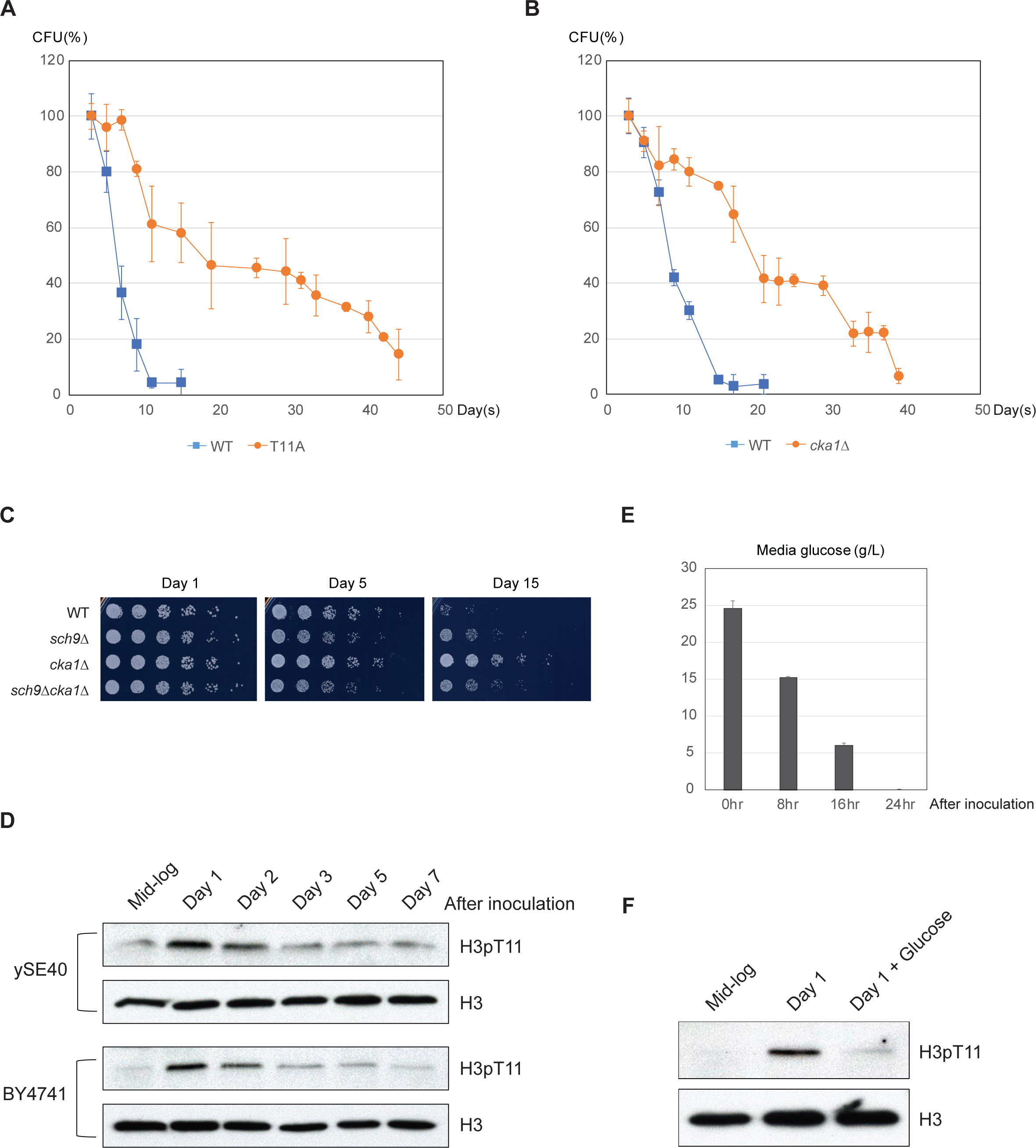
Phosphorylation of H3T11 regulates lifespan. (A) Chronological lifespan (CLS) assays of WT (ySE40) and H3T11A strains. (B) CLS assays of WT (BY4742) and *cka1Δ* strains. Error bars in CLS assays indicate STD from three to six biological replicates. CFU, colony forming units. (C) Relative viability of WT (BY4742), *sch9Δ, cka1Δ,* and *sch9Δcka1Δ* during CLS analysis at indicated time points. (D) H3pT11 levels measured at indicated times during CLS analysis of ySE40 (WT of histone mutant strains) and BY4741 strain analyzed by Western blots. (E) A Bar graph displaying media glucose concentration measured from the WT strain culture at indicated times in CLS analysis. Error bars indicate STD of three biological replicates. (F) H3pT11 levels of WT strain at exponential growth stage (mid-log), saturated day 1 culture (day 1), and day 1 culture with re-supplemented glucose (day 1 + glucose) analyzed by Western blots. For day 1 + glucose culture, 2% glucose was directly added to saturated day 1 culture, then incubated for additional 1 hour.

Sch9 and CK2 might regulate the lifespan by controlling the phosphorylation of H3T11 in response to alteration of glucose levels during chronological aging. To address this hypothesis, we tracked H3pT11 levels during the CLS assay. Interestingly, global levels of H3pT11 were significantly increased at day 1 after inoculation and then reduced (Figure 5D). At this time point, media glucose is depleted by consumption (Figure 5E), and yeast cells begin to change the utilization of its carbon source metabolism from fermentation to respiration (DeRisi et al., 1997; Galdieri et al., 2010), the process regulated by H3pT11 in nutritional stress conditions (Figure 2). Supplying glucose at day1 after inoculation suppressed the elevation of H3pT11 levels (Figure 5F). These data suggest that the increase in H3pT11 levels at early stage of CLS is anti-correlated with glucose availability in the media, and H3T11 phosphorylation mediated by Sch9 and CK2 affects lifespan by regulating the metabolic transition at this time point.

### H3pT11 controls lifespan by regulation of acid stress response

Upon depletion of glucose, yeast cells encounter several stresses including media acidification. Glucose depleted media becomes acidified, especially by acetic acid produced during early stage of the CLS assay. Media acidification has been suggested as a pro-aging factor, and the glucose sensing pathway via Sch9 is responsible for acetic acid stress response (Burtner, Murakami, Kennedy, & Kaeberlein, 2009; Fabrizio et al., 2005; Longo et al., 2012). To know whether impaired H3pT11 affects acidification or the levels of acetic acid in the media, we measured media acetate level and pH during CLS analysis. There were no significant differences in media pH and acetate level between WT and H3T11A during the first few days of CLS (Figure 6-figure supplement 1A and 1B). Indeed, the media acetate levels were even slightly higher in H3T11A mutant. However, H3T11A, and *cka1Δ* mutants, as well as *sch9Δ* mutant displayed strong resistance against high concentration of acetic acid in the media (Figures 6A and 6B). In addition, *sch9Δcka1Δ* mutant showed similar acetic acid resistance to *sch9Δ* single mutant (Figure 6B), as in the case of lifespan control (Figure 5C). We thought that extended lifespan in H3T11 phosphorylation defective mutants (H3T11A, *cka1Δ, sch9Δ*, and *sch9Δcka1Δ*) might be correlated to their resistance to acetic acid. Supporting this idea, buffering media to pH 6.0 abolished the extension of lifespan in H3T11A or *cka1Δ* (Figure 6C and Figure 6-figure supplement 1C). High level of acetic acid in the media disrupts glucose metabolism (Sousa, Ludovico, Rodrigues, Leão, & Côrte-Real, 2012). We observed that media glucose levels remained stable (i.e. glucose was not consumed) even after 24 hours of WT culture in 50 mM acetic acid containing SDC media (Figure 6-figure supplement 1D), suggesting impairment of glucose utilization after acetic acid treatment. By contrast, media glucose was consumed in 10 mM acetic acid condition, similar to the physiological acetic acid concentration of the media during CLS analysis (Figure 6-figure supplement 1D)(Longo et al., 2012). As H3pT11 responds to low glucose condition (Figure 1), we tested whether media acetic acid affects H3pT11 level. Indeed, H3pT11 was induced by 50mM acetic acid treatment, but not by 10mM acetic acid treatment (Figure 6-figure supplement 1E). In 50mM acetic acid condition, the H3pT11 increase was impaired in *cka1Δ, sch9Δ,* and *cka1Δsch9Δ* mutants (Figures 6D and 6E). These data indicate that acetic acid and low glucose stress response participate in the same pathway depending on both Sch9 and CK2. Hence, we conclude that H3pT11 mediated by CK2 and Sch9 controls lifespan by regulation of the stress responses at early stage of CLS.

**Figure 6.**
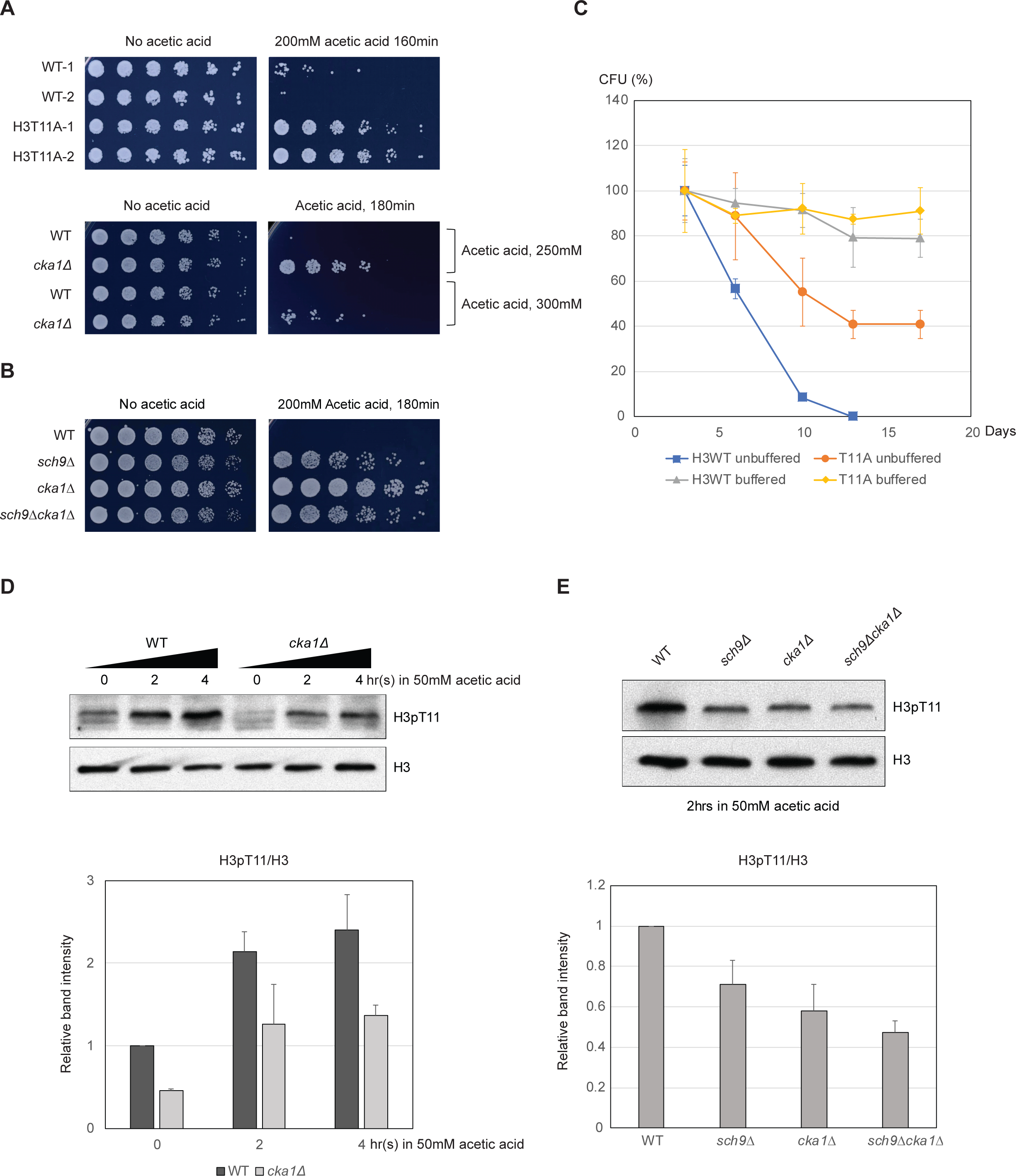
H3pT11 affects lifespan by regulation of acetic acid resistance. (A) Relative viability of H3T11A and *cka1Δ* mutants compared to their WT strains after exposure to indicated durations and concentrations of acetic acid. (B) Acetic acid resistance of WT, *sch9Δ, cka1Δ,* and *sch9Δcka1Δ*. (C) CLS assays of WT and H3T11A strains in buffered (pH 6.0) or unbuffered conditions. (D) H3pT11 levels in WT and *cka1Δ* upon 50mM acetic acid addition analyzed by Western blots (upper). The relative band intensities of H3pT11 to H3 signals (lower). (E) H3pT11 levels in WT, *sch9Δ, cka1Δ,* and *sch9Δcka1Δ* at 2 hours after 50mM acetic acid treatment analyzed by Western blots (upper). The relative ratios of H3pT11 to H3 signals (lower). All error bars indicate STD from three biological replicates.

### H3T11 phosphorylation is increased in aged cells

Since H3pT11 defective mutants displayed extension of CLS, we asked if H3pT11 level was changed in aged cells. To address this question, we used the yeast Mother Cell Enrichment (MEP) system strain, which selects the mother cells in the presence of estradiol (D. L. Lindstrom & D. E. Gottschling, 2009) (Figure 7-figure supplement 1A). As expected, estradiol treated yeast populations accumulated bud scars, a sign of many iterated divisions, relative to no estradiol controls (Figure 7A), indicating this method selectively isolates aged cells. In these cells, we observed an increase in the levels of H4K16 acetylation (H4K16ac), which is well known marker for aged cells (Figure 7B) (W. Dang et al., 2009). Importantly, we found that H3pT11 levels were also significantly increased in the aged cells (Figure 7B). Addition of estradiol into the media or ectopic expression of Cre-EBD78 protein did not cause the increase of H3pT11 and H4K16ac (Figure 7-figure supplement 1B). Thus, the levels of H3T11 phosphorylation increased by aging.

**Figure 7.**
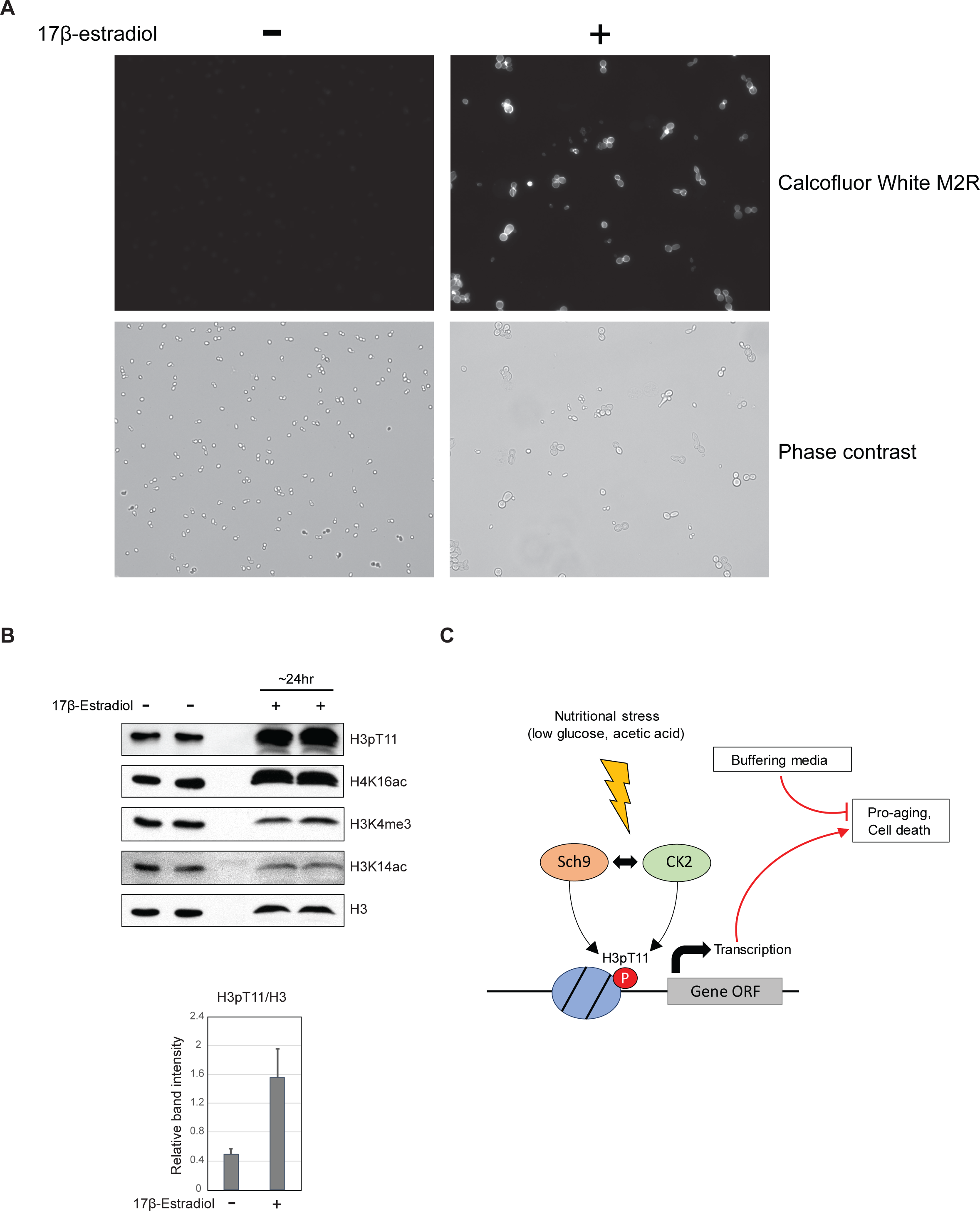
H3pT11 increases in aged cells. (A) Bud scar staining of MEP strain UCC8774 incubated with or without addition of 17β-estradiol. (B) (Left) H3pT11 levels of UCC8774 strain with or without addition of 17β-estradiol analyzed by Western blots. (bottom) The relative band intensities of H3pT11 to H3 signals from three biological replicates. Error bars represent STD. (C) Summary models of H3pT11 functions upon stress conditions.

## Discussion

### Cooperation of Sch9 and CK2

In this work, we identified novel functions of H3pT11 in nutritional stress conditions. H3pT11 specifically increased in nutritional stress conditions. H3pT11 regulates transcription upon stress and accelerates aging or cell death. The phosphorylation of H3T11 upon nutritional stress depends on the Sch9 and CK2 kinases (Summarized in Figure 7C). Our data indicate that Sch9 and CK2 act in overlapping manner in the regulation of H3pT11, acetic acid resistance, and aging (Figures 4C, 5C and 6B). Interestingly, CK2 has been shown to phosphorylate Sch9 human homologs S6 kinase (Panasyuk et al., 2006) and Akt1 (Di Maira et al., 2005), and Akt1 also phosphorylates CK2 (Mitchell, 2013). In addition, Sch9 and CK2 share some common targets in yeast, including transcription factor Maf1 and Bdp1 (Graczyk et al., 2011; Lee, Moir, & Willis, 2009, 2015). Sch9 has been suggested to bind to Ckb1, one of CK2 regulatory subunits (Fasolo et al., 2011). These data strongly suggest the intimate relationship between Sch9 and CK2. However, how these two enzymes coordinately regulate H3pT11 upon nutrient stress condition in yeast need to be determined.

### Transcription regulation by H3pT11

H3pT11 levels have been correlated with transcription regulation. In a human prostate tumor cell line, H3pT11 mediated by PKN1 is required for androgen stimulated gene transcription by facilitating removal of the repressive H3K9 methylation mark (Metzger et al., 2008). In mouse MEF cells, H3pT11 by Chk1 kinase is reduced upon DNA damage, and H3pT11 decrease causes repression of gene transcription, and reduction of H3K9 acetylation by GCN5 (Shimada et al., 2008). We also demonstrated the roles of H3pT11 in transcription regulation (Figure 2). Yeast Gcn5 has been suggested to bind better to H3S10 phosphorylation (H3pS10) or H3pT11 containing peptides than unmodified H3 peptides (Shimada et al., 2008). Interestingly, the study of Tetrahymena Gcn5 structure suggests that H3pS10 may facilitate the interaction between H3pT11 and Gcn5. It has been shown that H3pT11 is required for optimal transcription of H3pS10 dependent gene transcription in yeast (Clements et al., 2003), suggesting that crosstalk among H3pT11 and other histone modifications may have roles in the regulation of transcription.

### Lifespan regulation by H3pT11

H3pT11 defective mutants showed extended lifespan (Figure 5A) by altering stress response to low levels of glucose (Figure 2) and high levels of acetic acid (Figure 6A). High levels of acetic acid strongly disrupted glucose metabolism (Figure 6-figure supplement 1D) and induced H3pT11 (Figure 6C). Although we cannot exclude the possibility that other unknown stimuli are responsible for H3pT11 increase in acetic acid stress other than glucose starvation, the pathways regulated by Sch9 and CK2 were required for achieving acetic acid resistance and H3pT11 elevation (Figure 6D) in the case of low glucose (Figure 1). It has been suggested that utilizing ethanol and acetate, which are representative end-products of yeast fermentation, accelerates yeast aging, while maintaining glucose dependent pathways by gluconeogenesis is beneficial to yeast survival (Orlandi, Ronzulli, Casatta, & Vai, 2013). Recently, calorie restriction has been suggested to prolong lifespan by expanding the period of fermentation to respiration metabolic transition (Wierman, Maqani, Strickler, Li, & Smith, 2017). These results emphasize that the metabolic changes happened at early stage of CLS are critical for determination of lifespan, and our data indicate that H3pT11, mediated by Sch9 and CK2, is involved in the metabolic changes of this stage.

### H3pT11 upon aging

The levels of H3pT11 were increased when media glucose was depleted (Figures 1A and 1B). This feature is correlated with lifespan extension in H3pT11 defective mutants. The culture of aged MEP strain cells was not saturated in our experimental conditions (see STAR Methods), suggesting that the culture media contains sufficient glucose for their survival. It can be speculated that the metabolic states of aged cells resemble the nutritional stress state. In aged cells, the abundance of histones is decreased, and overexpression of histones extends lifespan in yeast (Feser et al., 2010; Hu et al., 2014; O’sullivan, Kubicek, Schreiber, & Karlseder, 2010). Interestingly, reduced histone abundance induces respiration in yeast (Galdieri, Zhang, Rogerson, & Vancura, 2016). Leaky induction of respiration by reduction of histone abundance may increase the levels of H3pT11 in aged cells. Comparison of metabolites in the cells between nutritional stress and aging would be useful for examining this.

H4K16ac is increased only in aged cells but not in nutritional stress condition (Figures 1A, 1B, and 7B). Thus, H4K16ac and H3pT11 may act in different pathways or in different periods during aging although both modifications are involved in lifespan regulation. The chromatin state in aged cells may be more complex, one of which may be similar to nutritional stress conditions. However, our study clearly showed the functional crosstalk between nutrition sensing pathways and chromatin regulation mediated by Sch9, CK2 and H3pT11, in controlling cellular lifespan.

## Materials and Methods

### Yeast strains

All yeast strains used in this study are described in Supplementary file 1. All single deletion mutants using KanMX4 marker and TAP tagged strains using HIS3 marker derived from BY4741 and BY4742 were obtained from Open Biosystems library (maintained at the Stowers Institute Molecular Biology facility). Histone H3 mutant shuffle strains were generated and maintained by Stowers Institute Molecular Biology facility (Nakanishi et al., 2008). Further deletion from these strains were achieved by targeted homologous recombination of PCR fragments containing marker genes flanked by the ends of the targeted genes. These strains were confirmed by PCR with primer set specific for their deletion marker or coding regions.

### Yeast culture conditions

Overnight saturated cell cultures were diluted into fresh YPD media and incubated until early mid-log phase. For nutritional stress experiments, these cultures were pelleted and washed once with YP containing no carbon source. Washed pellets were resuspended with YP media containing various concentrations of glucose as described in Figure 1A or 3% glycerol in elsewhere, and incubated at 30°C. For ‘glucose added’ samples in Figure 1-figure supplement 2A and Figure 5F, 2% glucose were directly added to the culture in YPglycerol (Figure 1-figure supplement 2A) or day 1 culture in SDC (Figure 5F) for 1hr, at 30°C. For *cdc19-1* temperature sensitive mutant in Figure S3A, the cells were cultured in YPD at 25°C and were then shifted to YP-glycerol media at 37°C.

### Isolation of aged cells using MEP strains

Isolation of aged cells using Mother cell Enrichment Program (MEP) strains was performed as previously described with minor modifications (Derek L Lindstrom & Daniel E Gottschling, 2009). Saturated cultures of MEP strains were diluted into 50 mL fresh YPD media and incubated at 30°C for overnight. Optical densities (OD) of the cultured cells were not exceed 1.0 after incubation. Cultures were inoculated to 100mL YPD media containing 1 μM 17β-Estradiol (Sigma) to a cell density of 4×10^3^ /mL to 4×10^4^ /mL. The cells were incubated at 30°C for 12 to 48 hours. Cell cultures were not saturated before preparation.

### Yeast bud scar staining

Cultured yeasts were fixed using 3.7% formaldehyde at 30°C for 30 minutes. 1×10^7^ cells were resuspended in 200 μL distilled water and were stained with 0.1mg/mL Fluorescent Brightener 28 (Sigma) at 30°C for 30 minutes. Stained cells were washed twice with 200μL water and were then resuspended in 50 μL water for imaging. The 10 μL of suspended cells were used for imaging with DAPI filter using fluorescent microscopy.

### Preparation of yeast whole cell extracts

Yeast whole cell extracts were prepared as previously described with minor modifications (Li et al., 2015). 5 OD cells were taken from 10 to 15 mL cultures. Harvested cell pellets were transferred to 1.7 mL Eppendorf tubes and washed once with 1 mL distilled water. Cell pellets were resuspended in 250 μL of 2M NaOH with 8% β-Mercaptoethanol and incubated on ice for 5 minutes. Cells were pelleted and washed once with 250 μL TAP extraction buffer (40 mM HEPES pH 7.5, 10% Glycerol, 350 mM NaCl, 0.1% Tween-20, phosphatase inhibitor cocktail from Roche, and proteinase inhibitor cocktail from Roche). Pelleted cells were resuspended in 180 μL 2X SDS sample buffer and boiled at 100°C for 5 minutes. 10 μL of each sample was used per lane for Western blotting.

### Chronological life span (CLS) assay

Chronological life span assay was performed as suggested previously with minor modifications (Longo et al., 2012). Saturated cultures in SDC media were diluted into fresh unbuffered SDC media or SDC media buffered at pH 6.0 by citrate-phosphate buffer (Burtner et al., 2009). Cultures were incubated at 30°C with 220 rpm shaking for aeration. At indicated times, same number of cells, based on optical density, were taken and plated into fresh YPD plate. The grown colony numbers were counted 2 days after the plating. Colony Forming Units (CFUs) were calculated by dividing the number of colonies grown at each time point by the number of colonies at day 3 (set as 100%).

### Chromatin IP

Chromatin IP assays were performed as previously described with minor modifications (Shim et al., 2012). 100 mL cultures were subjected to crosslinking by addition of 3 mL of 37% formaldehyde (Sigma) at RT for 15 minutes with constant swirling. 6mL of 2.5M glycine was added to quench crosslinking reaction at RT for 5 minutes. 80 OD quenched cells were pelleted by centrifugation at 6000 rpm for 5minutes and were washed twice with ice-cold 1X TBS (20 mM Tris pH 7.5 and 150 mM NaCl). Cells were lysed by bead beating in lysis buffer (50 mM HEPES pH 7.5, 150 mM NaCl, 1 mM EDTA, 1% Triton X-100, 0.1% Sodium deoxycholate, and 0.2% SDS). Lysates were sonicated to generate short DNA fragments using a Sonic Dismembrator Model 500 (Fisher) and were then clarified by centrifugation at 12000 rpm, 4°C for 20 minutes. Clarified lysates were diluted in four times volume of lysis buffer without SDS containing fresh Complete Mini protease inhibitor cocktail (Roche) and were then subjected to immunoprecipitation with antibodies against 4μL of H3pT11 (ab5168, Abcam) or 2μL of H3 (ab1719, Abcam). Antibody bound DNAs were recovered by incubation with 40 μL protein A agarose beads (GE Healthcare) at 4°C for overnight. Beads were washed sequentially with following buffers: once with lysis buffer without SDS for 10 minutes, twice with 500mM NaCl lysis buffer without SDS for 10minutes, once with LiCl buffer (10 mM Tris-HCl pH 8.0, 250 mM LiCl, 1 mM EDTA, 0.5% NP-40, and 0.5% Sodium deoxycholate) for 10minutes, and twice with TE buffer (10 mM Tris-Cl pH 7.5 and 1 mM EDTA) for 5minutes. The DNA/chromatin complexes were then eluted twice by incubation in elution buffer (1% SDS and 250 mM NaCl) at 65°C for 30 minutes with occasional vortexing. Elutes were treated with Proteinase K (Sigma) at 55°C for 2 hours and were then incubated at 65°C for overnight to reverse crosslinking. DNAs were prepared by phenol/chloroform extraction followed by ethanol precipitation. Precipitated DNAs were used for RT-qPCR or making libraries for ChIP-Sequencing.

### RNA purification

Yeast RNAs were prepared as previously described (Schmitt, Brown, & Trumpower, 1990). Briefly, 5 OD yeast cells were taken from cultures and were washed once with 1 mL of DEPC treated water. Washed pellets were transferred into 1.7 mL Eppendorf tubes and resuspended in 400 μL of AE buffer (50 mM sodium acetate pH5.3 and 10 mM EDTA). 40μl of 10% SDS was added to AE buffer resuspended cells and vortexed. 440 μL of phenol pH 8.0 (Sigma) was added to tubes, and then tubes were incubated at 65°C for 4 minutes. Tubes were rapidly cooled down in pre-chilled ice block until phenol crystals appear and were then centrifuged at 11000 rpm for 2minutes. Aqueous phase was carefully transferred into new tubes. RNAs in the aqueous phase were prepared using phenol/chloroform extraction followed by ethanol precipitation.

### TAP purification

TAP purification for CK2 complex was carried out as previously described (Li et al., 2015). 6L cultures of Cka1-TAP strain were grown in YPD medium at 30°C to an OD about 2.0 at 600 nm. The cell pellets were resuspended in TAP extraction buffer (40 mM HEPES pH 7.5, 10% Glycerol, 350 mM NaCl, 0.1% Tween-20, and protease inhibitor cocktail from Roche) and were then disrupted by bead beating. The crude cell extracts were treated with 125 U Benzonase and 50 μL of 10 mg/ml heparin at RT for 15 minutes to remove nucleic acid contamination and were then clarified by ultracentrifugation. Clarified extracts were incubated with IgG Sepharose (GE healthcare) beads at 4°C for 3 hours. The IgG-beads bound proteins were resuspended in TEV cleavage buffer (10 mM Tris pH 8.0, 150 mM NaCl, 0.1% NP-40, 0.5 mM EDTA, 10% glycerol, and Complete Mini protease inhibitor cocktail from Roche) and cleaved by addition of 5 μl of AcTEV (Invitrogen) at 4°C for overnight. The cleaved proteins were resuspended in calmodulin binding buffer (10 mM Tris pH 8.0, 300 mM NaCl, 1 mM magnesium acetate, 1 mM imidazole, 2 mM CaCl_2_, 0.1% NP-40, and 10% glycerol) and incubated with Calmodulin Sepharose (GE healthcare) beads at 4°C for 4 hours. Calmodulin-resin bound proteins were eluted by resuspension with calmodulin elution buffer (10 mM Tris pH 8.0, 150 mM NaCl, 1 mM magnesium acetate, 1 mM imidazole, 2 mM EGTA, 0.1% NP-40, 10% glycerol, and Complete Mini protease inhibitor cocktail from Roche).

### In vitro kinase assay

10 μL of TAP purified CK2 complex or Sch9 were incubated with 800 ng of recombinant Xenopus histone H3 with or without addition of 10 mM ATP in NEBuffer for protein kinase (NEB) at 30°C for 1 to 3 hours. The reactions were quenched by addition of SDS sample buffer and boiled at 100°C for 5 minutes.

### Spotting assay for acetic acid resistance

Overnight yeast cultures were diluted and were grown until their optical densities at 600 nm reach at mid-log phase. The cultures were treated with 200-300 mM acetic acid for 160-180 minutes. After the treatment, 4 folds serially diluted cells were spotted onto YPD plates. Plates were incubated at 30°C for 1 to 2 days for taking pictures.

### Media glucose and acetate quantification

Aliquots of yeast cultures were pelleted at indicated times, then supernatants were collected and frozen at −80°C until used. Glucose, and acetate concentrations in the growth medium were measured using enzymatic assay kits (EIAGLUC from ThermoFisher Scientific for glucose, and MAK086 from Sigma for acetate detection.) following manufacturer’s protocols.

### ChIP-sequencing and RNA-sequencing

ChIP-seq samples were sequenced in two lanes of an Illumina HiSeq 2500 at 51 bases, single end. Data was converted to fastq and demultiplexed using bcl2fastq. Reads were aligned to UCSC genome sacCer3 using bowtie2 (2.2.0) with option “-k 1”. Downstream analysis was done in R (3.2.2). Peaks were called using a custom perl script, requiring a 2-fold change between ip and input samples extending for 50 bases. Peaks closer than 400 bases were merged. Differential peaks of H3pT11 between glucose and glycerol were called using R package DiffBind (2.0.9). Genes closest to differential peaks were identified using bedtools closest (2.26.0) to identify the closest transcription start site. After removing any Pol III, tRNA, and rRNA genes, 366 peaks were found up in glycerol versus glucose, and 139 peaks were down in glycerol versus glucose. Gene ontology enrichment was performed using a hypergeometric test in R. Terms shown in the barplot had p-value < .05. The length of the bar represents the fold enrichment of the term’s frequency in the list given the frequency of the term in the genome. Metagene plots were generated in R using 101-base mean-smoothed windows +/− 2000 bases around the TSS or +/− 500 bases around the transcript region (start to end). Different length genes were accommodated using the approx() function, which uses linear interpolation to define the approximated data points. After getting approximated values for each gene, the mean value at each position was used to generate the plot.

RNA-seq samples were sequenced in two lanes of an Illumina HiSeq 2500 at 51 bases, single end. Data was converted to fastq and demultiplexed using bcl2fastq. Reads were aligned to UCSC genome sacCer3 with annotation from Ensembl 84 using tophat (2.0.13) with “-x 1 -g 1”. Reads were counted on genes (unioned exon space) using bedtools coverage (2.26.0). Data was read into R (3.2.2). Differentially expressed genes were found using R package edgeR (3.14.0) and required to have BH-adjusted p-value < .05 and two-fold change in order to be called differentially expressed. All correlations shown were calculated using cor () function in R, which is Pearson correlation by default. Lists of genes for the cyto/mito boxplot were taken from previous work (Cheng et al., 2007).

## Additional files

Supplementary file 1 – Yeast strains used in this study

**Figure 1-figure supplement 1.**
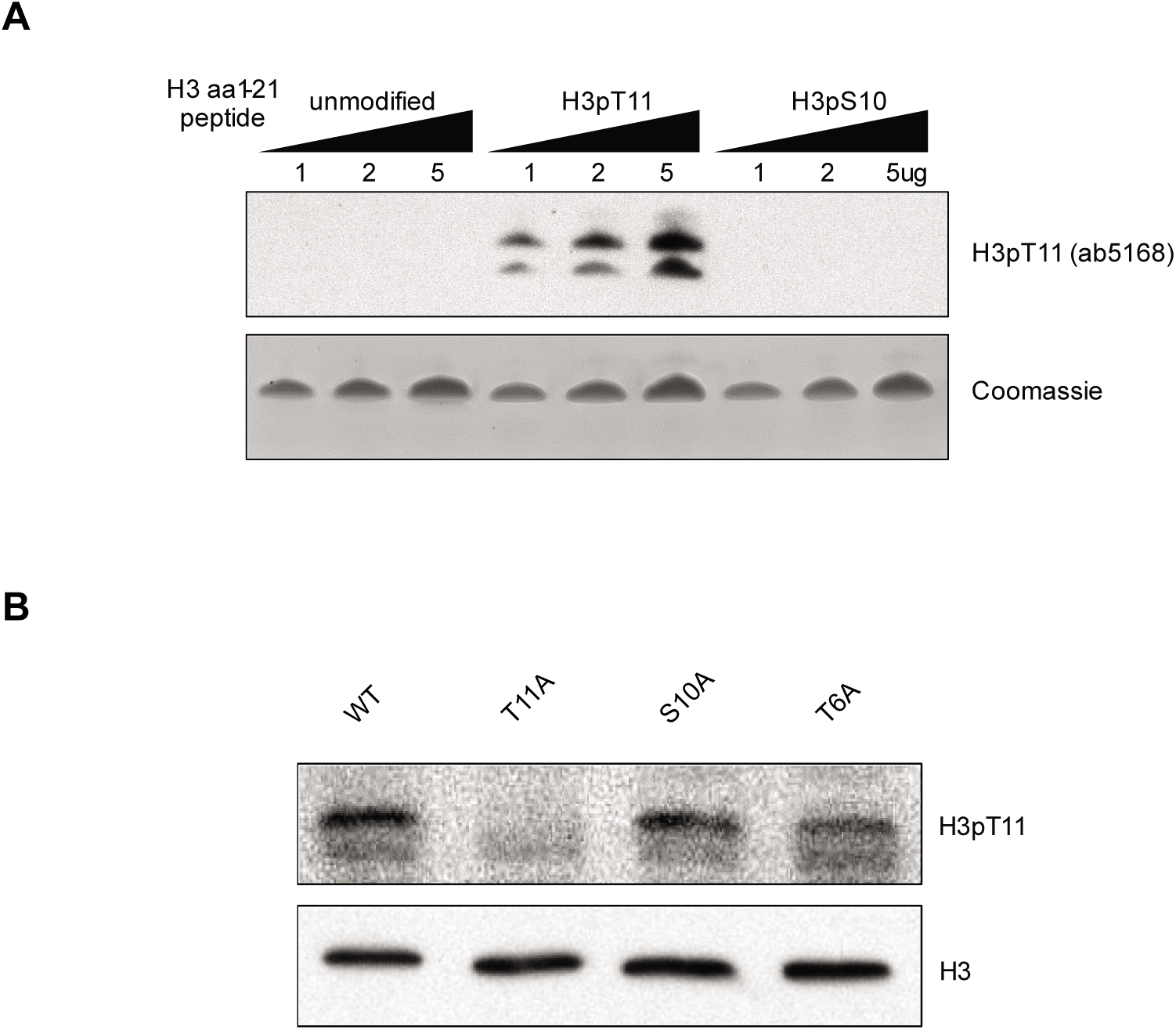
H3pT11 antibody validation. (A) H3pT11 antibody specificity test against unmodified, H3pT11, and H3pS10 containing histone H3 amino acids (aa) 1-21 peptides measured by Western blots. (B) H3pT11 levels in WT (ySE40), H3T6A, H3S10A and H3T11A strains in YPD condition.

**Figure 1-figure supplement 2.**
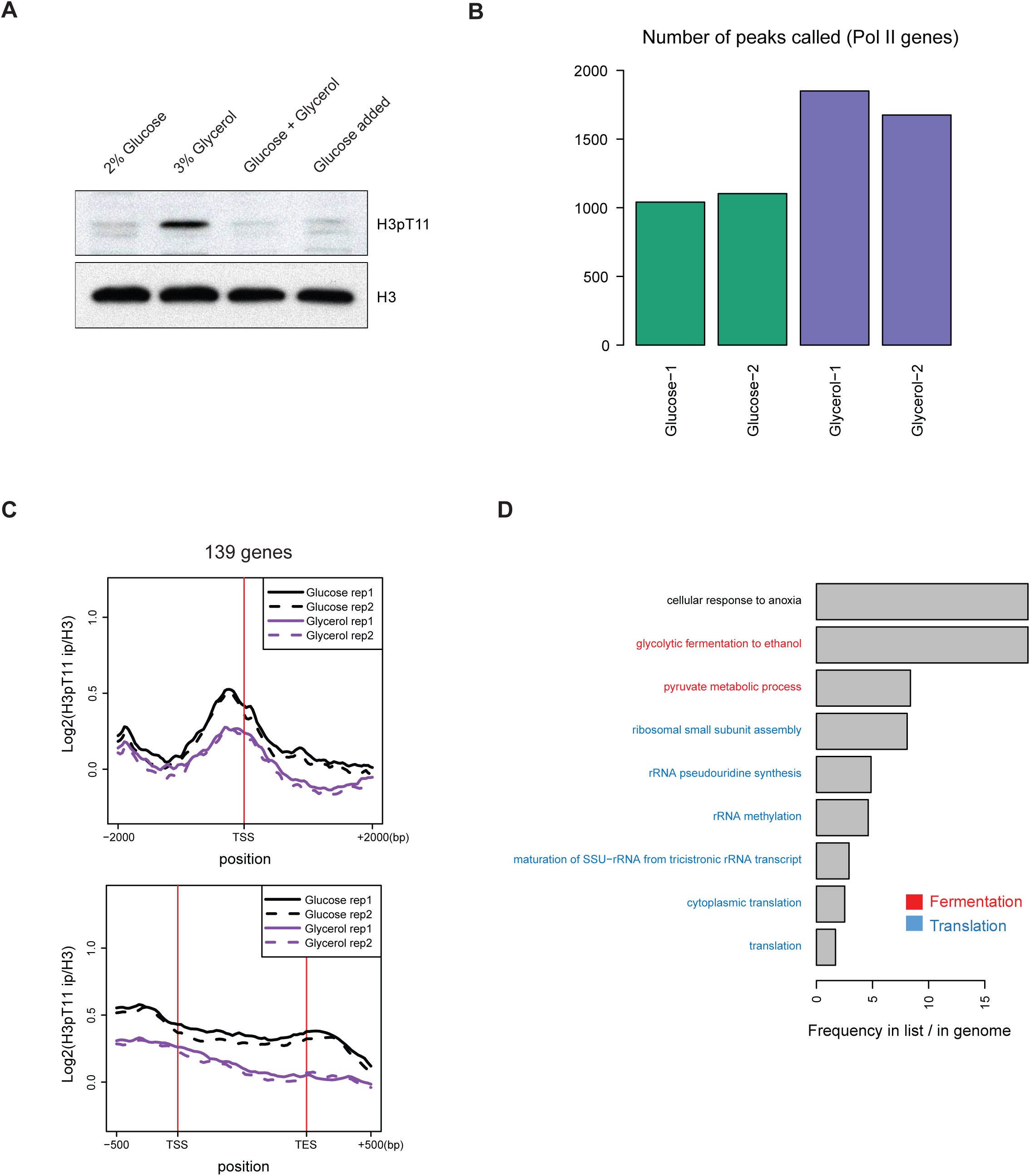
H3pT11 responds to low glucose condition. (A) Western blots showing H3pT11 levels in WT (BY4741) cultures shifted from YPD to YP media containing 2% glucose, 3% glycerol, or 2% glucose with 3% glycerol (Glucose + Glycerol) then incubated for 1 hour. For ‘Glucose added’ sample, WT cultures were initially shifted from YPD to YP with 3% glycerol for 1 hour, then 2% glucose was directly added to the culture, and further incubated for 1 hour. (B) A bar plot showing the total number of H3pT11 peaks in YPglycerol condition (Glycerol-1 and 2) and YPD condition (Glucose-1 and 2) ChIP-seq experiments. This bar plot does not include peaks overlapping non-pol II genes or peaks on the mitochondrial chromosome. (C) The average profiles of H3pT11 around the transcription start site (TSS) and across the gene body for 139 genes whose H3pT11 levels are decreased in YPglycerol (Glycerol) compared to YPD (Glucose) condition. tRNA and rRNA genes have been excluded. TES: transcription end site. (D) GO term analysis of the 139 genes from ChIP-sequencing data shown in Figure S2C. H3pT11 levels are decreased at genes related in fermentation (in red) and translation (in blue).

**Figure 3-figure supplement 1.**
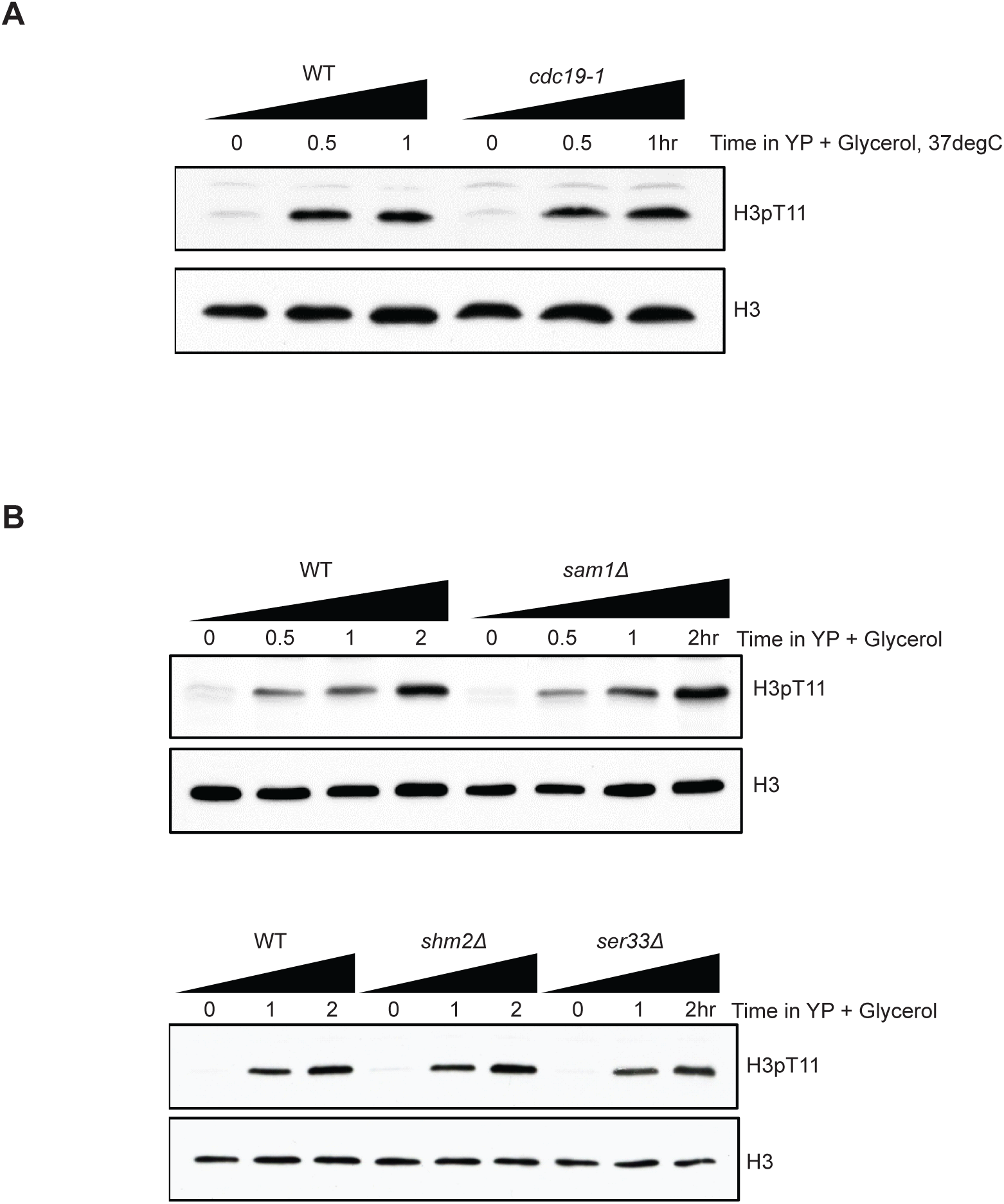
SESAME is not responsible for increased H3pT11 in nutritional stress condition. (A) Western blots showing that H3pT11 level changes of WT and pyruvate kinase 1 temperature sensitive mutant *cdc19-1* upon media shift from YPD, at permitting temperature (25°C) to YPglycerol, at non-permitting temperature (37°C). (B) H3pT11 levels in WT and SESAME subunit deletion mutants: *sam1Δ, shm2Δ*, and *ser33Δ* mutants upon media shift to YPglycerol analyzed by Western blots.

**Figure 3-figure supplement 2.** Cka1 is required for H3pT11. (A) H3pT11 levels in yeast kinase deletion mutants (*bub1Δ, cka1Δ*, and *cka2Δ*) and mutants of candidates for H3pT11 kinase (*mek1Δ* and *chk1Δ*) cultured in YPD media examined by western blots. (B) H3pT11 levels in subunits of CK2 deletion mutants: catalytic subunits, *cka1Δ* and *cka2Δ*, and regulatory subunits, *ckb1Δ* and *ckb2Δ,* cultured in YPD media examined by western blots.

**Figure 6-figure supplement 1.**
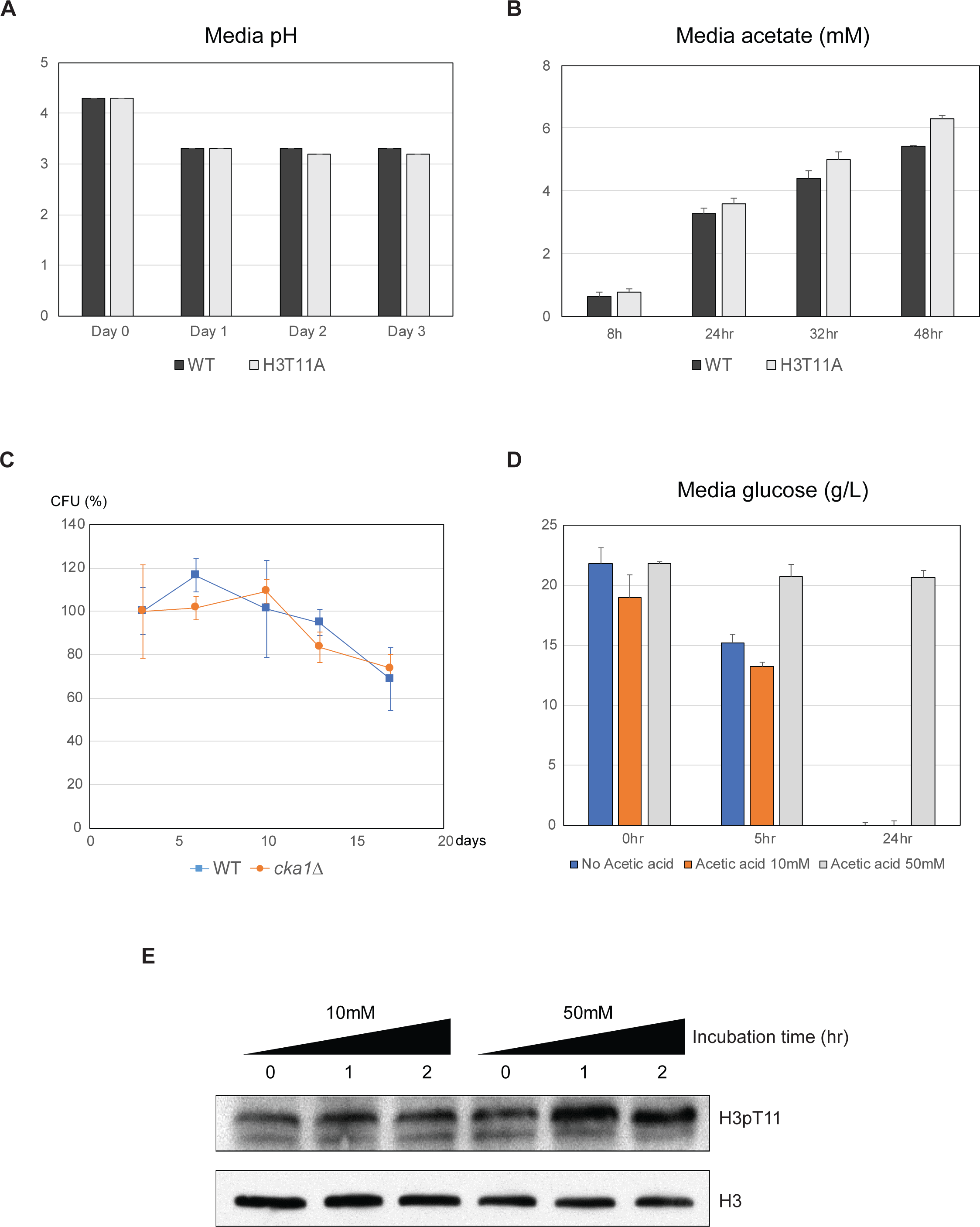
H3pT11 controls lifespan by regulation of acid stress response. (A) Media pH in WT cultures during CLS analysis. (B) Media acetate concentration in WT cultures at indicated times. (C) CLS analysis of WT and *cka1Δ* strain cultured in SDC media buffered at pH 6.0. (D) Media glucose concentrations of WT strain cultures in SDC media (no acetic acid) or SDC media supplemented with 10mM or 50mM acetic acid. Acetic acid treatment time was regarded as 0 hour. From (A) to (D), All error bars indicate standard deviation (STD) of three biological replicates. (E) H3pT11 levels in WT strain upon treatment of 10mM or 50mM acetic acid measured by Western blots.

**Figure 7-figure supplement 1.**
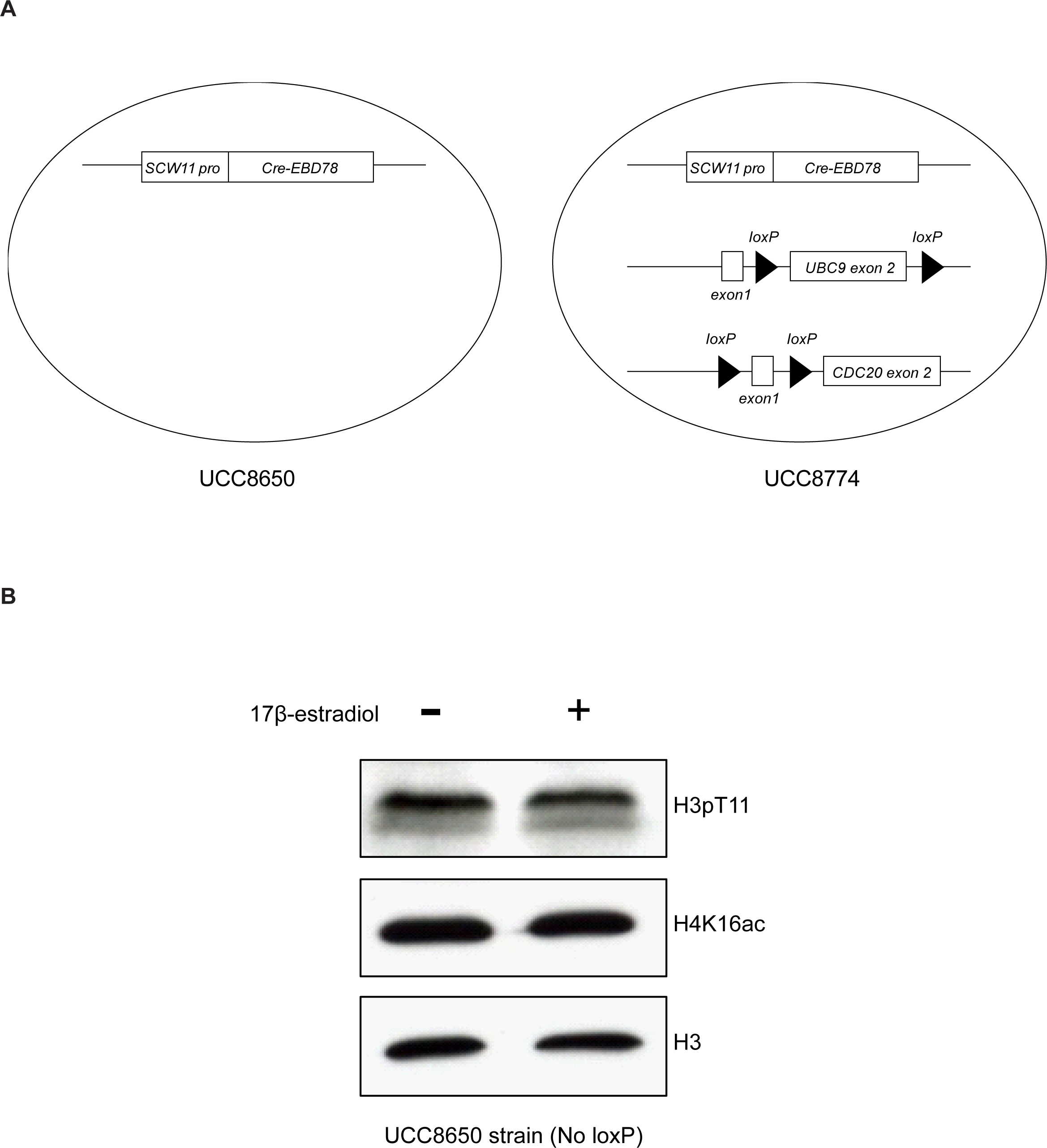
Strain of Mother cell Enrichment Program (MEP) for isolation of aged cells. (A) Schematic diagram of MEP strain. UCC8774 (Right) strain contains Cre-EBD78 gene construct governed by yeast Scw11 gene promoter (Scw11 pro), which is only active in recently budded daughter cells. UCC8774 strain also has loxP sequences surrounding exons of two yeast essential genes; UBC9 and CDC20, while UCC8650 strain (Left) does not. Consequently, UCC8774 daughter cells cannot survive in the presence of 17β-estradiol by removal of UBC9 and CDC20 exons by Cre recombinase, but UCC8650 strain can survive. (B) H3pT11 and H4K16ac levels of UCC8650 strain with or without addition of 17β-estradiol measured by Western blots. UCC8650 daughter can survive in the presence of 17β-estradiol while expressing Cre-EBD78.

